# GAIT-GM integrative cross-omics analyses reveal cholinergic defects in a *C. elegans* model of Parkinson’s disease

**DOI:** 10.1101/2021.07.16.452702

**Authors:** Danielle E. Mor, Francisco Huertas, Alison M. Morse, Rachel Kaletsky, Coleen T. Murphy, Vrinda Kalia, Gary W. Miller, Olexander Moskalenko, Ana Conesa, Lauren M. McIntyre

## Abstract

**Background:** Parkinson’s disease (PD) is a disabling neurodegenerative disorder in which multiple cell types, including dopaminergic and cholinergic neurons, are affected. The mechanisms of neurodegeneration in PD are unknown, limiting the development of therapies directed at disease-relevant molecular targets. *C. elegans* is a genetically tractable model system that can be used to disentangle disease mechanisms in complex diseases such as PD. Such mechanisms can be studied combining high-throughput molecular profiling technologies such as transcriptomics and metabolomics. However, the integrative analysis of multi-omics data in order to unravel disease mechanisms is a challenging task without advanced bioinformatics training. Galaxy, a widely-used resource for enabling bioinformatics analysis by the broad scientific community, has poor representation of multi-omics integration pipelines.

**Results:** We present the integrative analysis of gene expression and metabolite levels of a *C. elegans* PD model using GAIT-GM, a new Galaxy tool for multi-omics data analysis. Using GAIT-GM, we discovered an association between branched-chain amino acid metabolism and cholinergic neurons in the *C. elegans* PD model. An independent follow-up experiment uncovered cholinergic neurodegeneration in the *C. elegans* model that is consistent with cholinergic cell loss observed in PD.

**Conclusion:** GAIT-GM is an easy to use Galaxy-based tool for generating novel testable hypotheses of disease mechanisms involving gene-metabolite relationships.

## 1 Background

Parkinson’s disease (PD) is a progressive neurodegenerative disorder that is characterized by motor symptoms including resting tremor, muscle rigidity, and slowness of movement (Hoehn and Yahr, 1967), as well as non-motor symptoms including sleep disturbances and autonomic dysfunction (Lim et al., 2009). Several populations of neurons degenerate in PD, including dopaminergic neurons in the substantia nigra, and cholinergic neurons in the dorsal motor nucleus of the vagus and the nucleus basalis of Meynert (Forno, 1996). Current treatment options mitigate symptoms of the disease but do not offer a cure, emphasizing the urgent need for greater understanding of disease mechanisms to guide the design of new potential therapeutics.

*C. elegans* is a small nematode worm with an exceptionally high degree of genetic tractability that can be used to uncover biological mechanisms in a tissue and cell type-specific manner. *C. elegans* possess orthologs for 60-80% of human genes, and the worm nervous system utilizes many of the same neurotransmitters as in mammals, including dopamine, acetylcholine, γ-amino butyric acid (GABA), and glutamate (Corsi et al., 2015). In our recently developed *C. elegans* model of PD, RNAi-mediated reduction of the branched-chain amino acid (BCAA) transferase *bcat-1* resulted in PD-like motor dysfunction and dopamine neuron degeneration (Yao & Kaletsky et al., 2018). Transcriptomics and metabolomics profiling were used to study underlying molecular mechanisms, and analysis of each of these datasets separately pointed to an important role of mitochondria in driving disease (Mor et al., 2020; Mor & Murphy, 2020).

Integration of gene expression and metabolomics data requires advanced bioinformatics skills that are not always within the reach of the biologists who develop and test disease models. However, the experimental scientists are in an advantageous position to interpret the molecular data due to their profound knowledge of the biology of the system and of the significance of analysis results. Therefore, user-friendly bioinformatics tools that provide experimentalists with access to advanced omics data integration analysis methods are needed. Galaxy is a widely used open-source community development platform with an easy to understand GUI interface and a ‘mix and match’ pipeline building philosophy that makes it attractive to users with limited bioinformatic background. Galaxy has become one of the most successful resources for reproducible omics data analysis and already contains a plethora of tools to analyze gene expression, and metabolomics (e.g. Kirpich et al. 2018). However, tools for the integrative analysis of transcriptomics and metabolomics data are not readily available in Galaxy.

GAIT-GM is a new Galaxy tool developed to address these limitations. GAIT-GM leverages the plethora of tools available in Galaxy for the separate analysis of transcriptomics and metabolomics data to create an integration module that combines significant gene expression and metabolite annotations. This allows for studies of both data-driven and biologically oriented relationships among genes and metabolites, while seamlessly managing the data formatting issues that are endemic in trying to integrate these data types. GAIT-GM integration is based on the joint mapping of genes and metabolites to the KEGG database and in the application of dimension reduction (Ponzoni et al., 2014) and clustering (Stone and Ayroles, 2009) techniques to combine gene expression and metabolite level signals. In addition to this pathway-based integration, GAIT-GM includes unbiased approaches that directly explore links between metabolites and gene expression by modeling metabolite changes as a function of transcriptional regulation (Figure 1). We have applied these tools to the *bcat-1(RNAi) C. elegans* transcriptomics and metabolomics datasets in order to gain deeper insights into *bcat-1*-associated PD phenotypes. Using GAIT-GM, a previously underappreciated relationship between BCAA metabolism and cholinergic signaling were identified. This relationship was independently tested with new experimental data. These independent follow-up experiments with *C. elegans* confirmed that *bcat-1* reduction causes cholinergic neurodegeneration, paralleling the loss of cholinergic neurons in PD.

**Figure 1:**
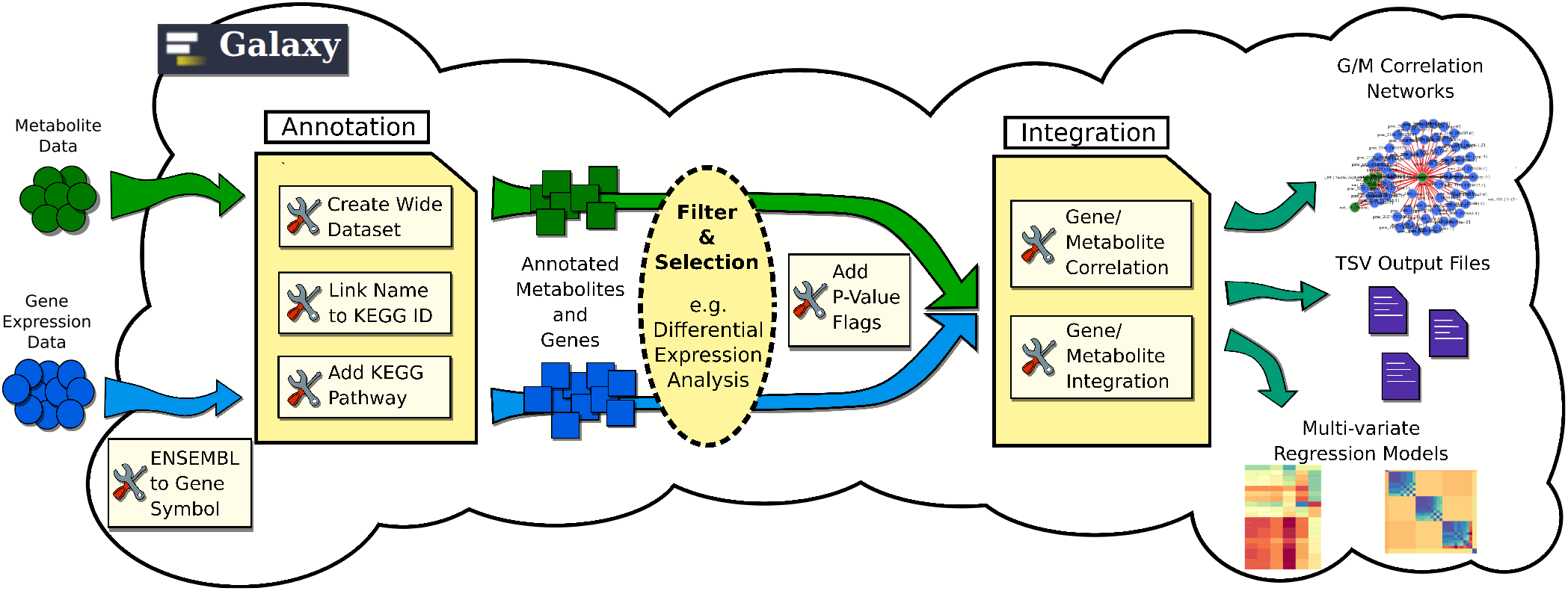
Galaxy pipeline construction enabled by GAIT-GM. Metabolomics and gene expression data are mapped to KEGG pathways. Relevant features are identified using existing statistical analysis approaches. Changes in metabolite levels are modeled as a result of changes in gene expression. Pipelines for data-driven and biology-informed integration are easily created.

## 2 Results

### 2.1 GAIT-GM Annotation Tool effectively maps genes and metabolites to pathways

Annotation using GAIT-GM text mining (see Supplementary Methods) resulted in 110/110 metabolites (100%) mapping to KEGG compound IDs and 3,096/3,146 genes (98%) mapping to KEGG IDs. While some gene and metabolite KEGG IDs did not have associated pathways, 79 metabolites were mapped to a total of 75 unique pathways, and 1187 genes mapped to a total of 115 unique pathways (Supplementary Figure 1). Pathways where both metabolites and genes were found included valine, leucine and isoleucine degradation; alanine, aspartate and glutamate metabolism; glycolysis / gluconeogenesis; and citrate cycle (TCA cycle), which recapitulates the pathway identification obtained at the separate analysis of these data (Mor et al., 2020).

### 2.2 GAIT-GM Integrative Tools characterize the gene-metabolite co-variation network

We first hypothesized that metabolic changes in *bcat-1(RNAi)* worms may relate to global transcriptional patterns. To explore this idea, we used MMC to cluster metabolites into modules and summarized pathway transcriptional activity by obtaining metagene profiles using PANA (see Methods). MMC identified nine metabolite modules (Figure 2A). Several of these modules consisted of metabolites related to BCAA metabolism, including L-valine (Module 6), succinyl-CoA (Module 3), and thiamin diphosphate (Module 1). sPLS analysis of Module 6 revealed a negative correlation between L-valine and the valine, leucine and isoleucine degradation pathway (Figure 2B), which is consistent with the known increase of BCAAs in the *bcat-1* background (Mansfeld et al., 2015; Mor et al., 2020). Also consistent with previous findings was the positive correlation between L-valine and the oxidative phosphorylation pathway, since *bcat-1* knockdown was reported to increase mitochondrial respiration (Mor et al., 2020). In addition, L-valine was positively correlated with the phenylalanine, tyrosine and tryptophan biosynthesis pathway and negatively correlated with homologous recombination, entirely consistent with previously identified upregulation and downregulation of these pathways, respectively, in *bcat-1(RNAi)* worms (Mor et al., 2020). In summary, integrative results involving L-valine recapitulated previous observations and provided confidence on the power of the GAIT-GM approach to recover meaningful biological insights.

**Figure 2:**
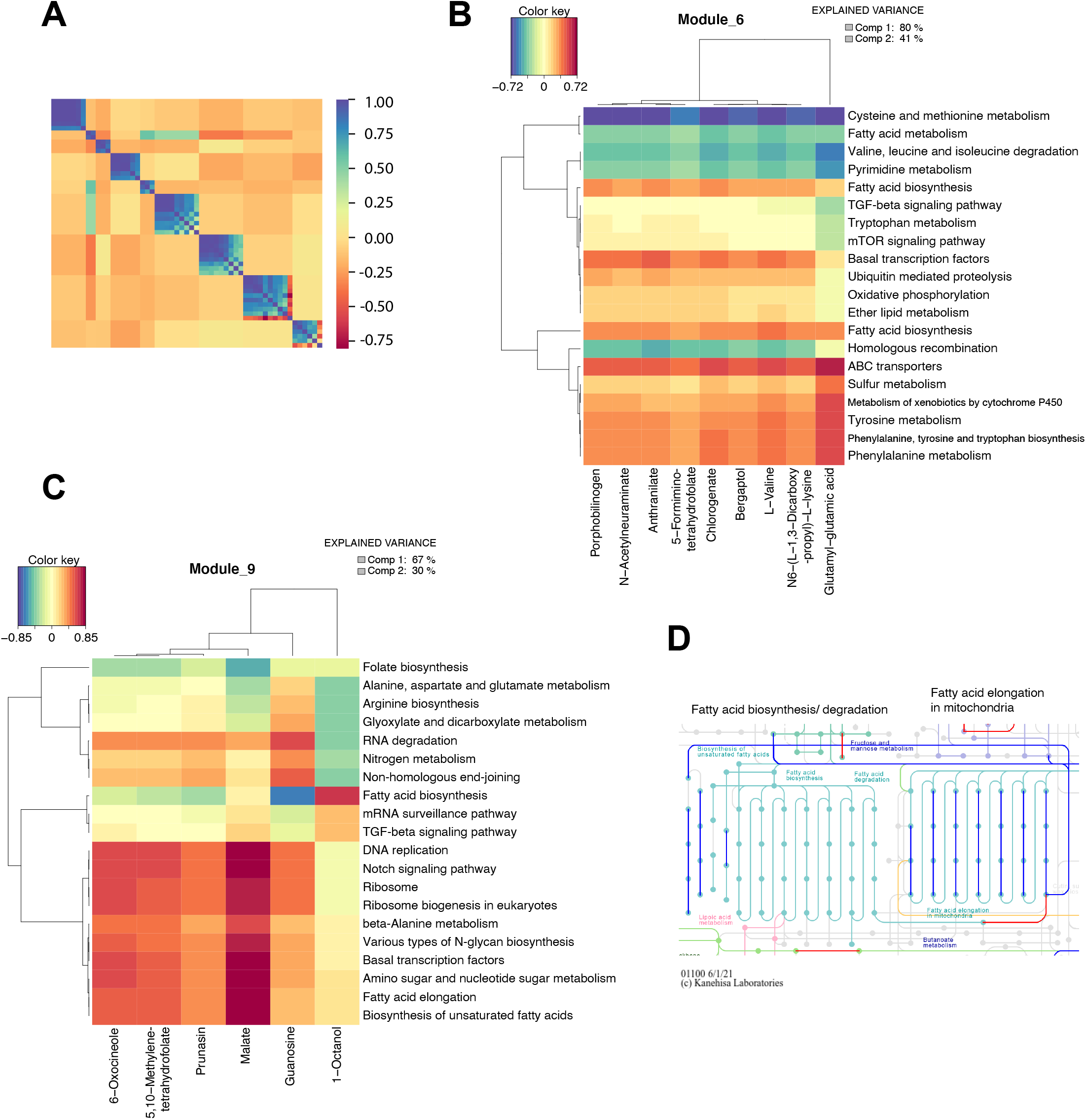
Integration analysis highlights gene-metabolite pathway relationships. **(A)** Modulated Modularity Clustering (MMC) analysis in which a smooth correlation ordering is performed in order to differentiate blocks of metabolites with the same behavior. Metabolites were grouped into 9 different modules. Blue is positively correlated and red is negatively correlated. **(B-C)** Heatmaps of Modules 6 **(B)** and 9 **(C)** after a sparse PLS (sPLS) analysis was performed with the Gene/Metabolite Integration Tool. Metagenes were estimated from the PANA approach. **(D)** Representations of *C. elegans* fatty acid metabolic pathways in KEGG. Genes significantly downregulated in *bcat-1(RNAi)* worms are blue, and genes significantly upregulated in *bcat-1(RNAi)* worms are red.

Module 9 included the TCA cycle metabolite, malate (Figure 2C), which is thought to be decreased upon *bcat-1* knockdown along with other TCA cycle substrates (Mor et al., 2020). In Module 9, malate was highly positively correlated with DNA replication and Notch signaling (Figure 2C), two pathways known to be downregulated in *bcat-1(RNAi)* worms (Mor et al., 2020), thus maintaining the expected directional relationship. Interestingly, both Modules 6 and 9 involved several pathways related to fatty acid metabolism, including fatty acid biosynthesis, fatty acid elongation, and biosynthesis of unsaturated fatty acids (Figures 2B, C). Closer examination of the fatty acid metabolic pathways revealed that the associated genes are largely downregulated, particularly those functioning in fatty acid elongation in mitochondria (Figure 2D). Given that mitochondrial respiration is increased in *bcat-1(RNAi)* worms (Mor et al., 2020), it possible that fatty acid elongation is being downregulated in favor of fatty acid breakdown and utilization as a substrate for TCA cycle activity. Fatty acid metabolism was not previously highlighted in single omics analyses of the *bcat-1(RNAi)* worms, indicating that GAIT-GM integrated analysis may be used to reveal new potential relationships within networks of co-regulated genes and metabolites.

### 2.3 GAIT-GM data-driven analysis points to potential cholinergic neuron defects

To explore possible gene-metabolite relationships not captured by the pathway analysis, we performed an unbiased discovery analysis using the Pearson correlation between individual differentially expressed metabolites and genes (Figure 3). Two prominent metabolite hubs with 226 and 60 highly correlated genes, respectively, were identified (Figure 3A). In the larger cluster of genes, which are positively correlated with the metabolites retinoate and amylopectin, several functional terms related to neurotransmission were significantly overrepresented. These include chemical synaptic transmission, neurotransmitter secretion, and synaptic vesicle cycle (Figure 3B). The known role of retinoic acid in neuronal development and maintenance (Maden, 2007) offers a possible explanation for the high number of neuronal genes correlated with this metabolite.

**Figure 3:**
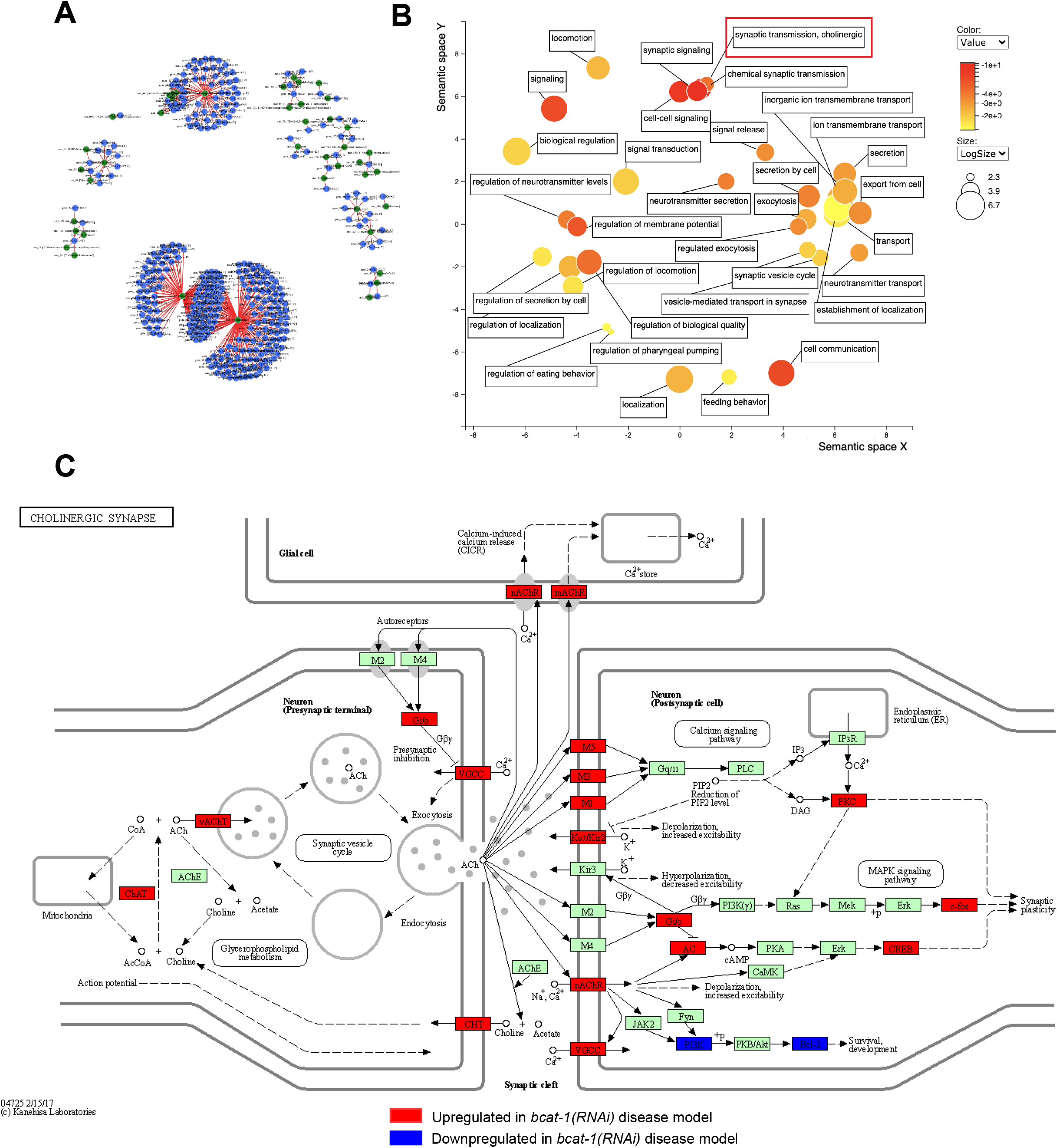
Correlation analysis between individual genes and metabolites points to potential cholinergic defects. **(A)** Output from the Gene/Metabolite Correlation Tool showing the 500 strongest correlations between individual genes and metabolites. Metabolites are represented in green, while genes are represented in blue. Red lines represent positive correlations and light blue lines signify negative correlations. **(B)** Gene Ontology analysis visualized by REVIGO for the largest cluster from **(A)** shows functional enrichment for neurotransmission, and in particular cholinergic synaptic transmission (red box). **(C)** Representation of the human cholinergic synapse KEGG pathway with *C. elegans* gene orthologs significantly upregulated in *bcat-1(RNAi)* worms in red, gene orthologs significantly downregulated in *bcat-1(RNAi)* worms in blue, and all other genes with KEGG annotations in green.

The second largest cluster was positively correlated with arachidonyltrifluoromethane, a derivative of arachidonic acid, which also plays a role in neuronal function (Bosetti, 2007). However, in contrast to the first cluster, genes in the second largest cluster were functionally enriched for processes related to reproduction, including germ cell development, germ-line stem cell division, and cellular process involved in reproduction in multicellular organism (Supplementary Figure 2). These findings suggest the interesting possibility that *bcat-1* disruption may affect reproduction in addition to neuronal health. Thus, GAIT-GM analyses offer several avenues by which to further investigate *bcat-1* regulation of biological and disease processes.

Here were focus on the impact of *bcat-1* on the regulation of neuronal function and its role in neurodegenerative disease. We identified a significant functional enrichment related to cholinergic synaptic transmission (Figure 3B) in GO analysis of the larger cluster. Further examination of cholinergic synapse components revealed that the associated genes are largely upregulated in *bcat-1(RNAi)* worms, including the vesicular acetylcholine transporter VACHT/*unc-17*, and the choline acetyltransferase CHAT/*cha-1* (Figure 3C). These findings suggested to us that *bcat-1(RNAi)* worms may have abnormal cholinergic signaling and/or damage to cholinergic neurons which may contribute to PD-like disease.

### 2.4 Follow-up confirmation of cholinergic degeneration in the *C. elegans* PD model

To investigate the possibility of cholinergic defects, neuronal RNAi-sensitive worms were fed *bcat-1* or control (empty vector) RNAi at the onset of adulthood until day 8, at which time cholinergic neurons expressing GFP were imaged. Remarkably, RNAi-mediated reduction of *bcat-1* resulted in significant degeneration of cholinergic neurites in the head, ventral nerve cord, and dorsal nerve cord (Figure 4). Specifically, cholinergic neurites in *bcat-1(RNAi)* worms had more severely degenerated morphologies than control RNAi-treated animals, including wavy appearances and defasciculation (Figure 4). These findings indicate that BCAA metabolism plays a crucial role in the health of cholinergic neurons and illustrate the power of GAIT-GM to integrate metabolite and gene expression data to reveal novel mechanisms of disease.

**Figure 4:**
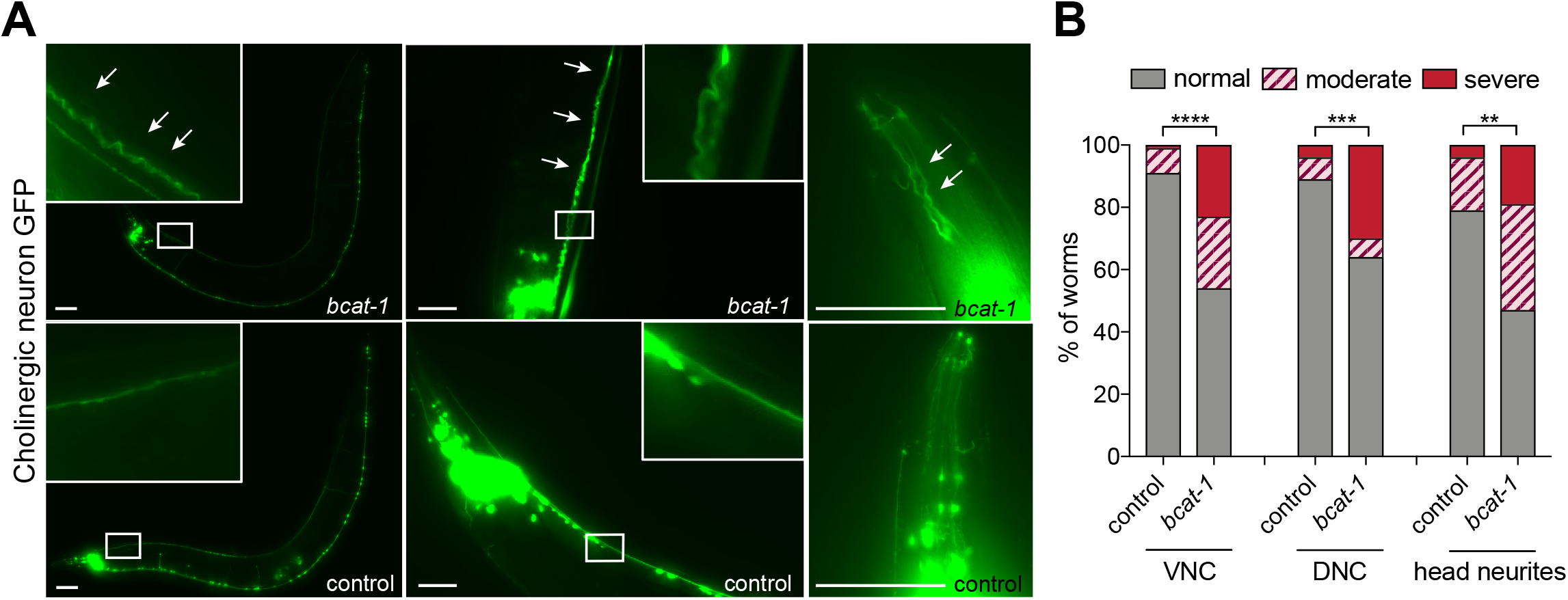
*bcat-1* RNAi-treated worms exhibit cholinergic neurodegeneration. **(A-B)** Cholinergic neuron projections in the head, ventral nerve cord (VNC), and dorsal nerve cord (DNC) of RNAi-sensitive worms (*unc119p::sid-1;unc-17p::GFP*) were scored for the degree of degenerated appearance on day 8 following *bcat-1* or control RNAi treatment during adulthood. In **(A)**, representative images of DNC (left), VNC (middle), and head neurites (right) are shown, with arrows indicating abnormal neurite regions. Scale bars, 50 μm. Quantification in **(B)**. Chi-square test in, n ≥ 60 worms per group. **p < 0.01; ***p <0.001, ****p <0.0001.

## 3 Discussion

Parkinson’s disease (PD) is increasingly recognized as a multi-system disorder involving multiple brain regions and neurotransmitter systems, yet the mechanisms causing neurodegeneration of various neuronal subtypes is largely unknown. The separate analysis of gene expression and metabolomics data in a *C. elegans* model of PD resulted in valuable new discoveries, in particular that *bcat-1* knockdown in worms increases mitochondrial respiration, and reducing mitochondrial activity with low-dose treatment of sodium azide mitigates PD-like motor dysfunction and dopaminergic neurodegeneration in *bcat-1(RNAi)* worms (Mor et al., 2020), underscoring the power of these technologies.

An integrative analysis of the transcriptomics and metabolomics data has the potential to reveal deeper insights into the interplay between gene expression and metabolic changes associated with PD. Existing methods for the integration of metabolomics and gene expression data are often based on multivariate dimension reduction techniques that inform on the covariation structures between transcript and metabolite levels (Cavil et al 2015, Meng et al 2016). These methods rarely incorporate existing knowledge, which hampers the interpretation of analysis results. Pathway databases such as KEGG, that include both metabolites and genes, can be used to map features to a common biological process and to guide integrative analysis and interpretation. However, mapping to pathways such as in KEGG is limited by the sparsity of metabolites captured in particular metabolic pathways. Conversely, many detected metabolites that cannot be mapped to the pathway data are eventually left aside in the integrative analysis. Moreover, despite the clear evidence of the importance of using metabolite identifiers (Kind, Scholz and Fiehn 2009), names are still prevalent, impeding the use of databases that rely on identifiers. Finally, existing pathway tools have limited or no choices for the preprocessing of gene expression and metabolomics data, thus fragmenting the analysis pipeline.

To meet these challenges, we developed GAIT-GM, a Galaxy-based tool for the integrative analysis of multi-omics data. Galaxy is a platform widely used by experimentalists who analyze their own omics data because it allows flexible configuration of analysis pipelines to accommodate a wide range of experimental designs and analysis needs. However, few tools for integrative analysis of transcriptomics and metabolomics datasets exist in Galaxy. The GAIT-GM tool fills a gap in multi-omics data analysis and enables both hypothesis-driven and unbiased hypothesis-generating integrated data analysis in a well-supported open source environment. The tool provides a user-friendly, modular analytical framework that enables the construction of reproducible pipelines and transparent sharing of analytical approaches and results. Additionally, there are two important yet unmet needs in the analysis of metabolomics data specifically developed here: the efficient mapping of compound names to KEGG, and the analysis of the heterogeneity and co-regulation with gene expression to improve functional characterization.

Here, unbiased analysis of the association between metabolites and gene expression in *bcat-1(RNAi)* versus control *C. elegans* using GAIT-GM led to the discovery of cholinergic neurons as a vulnerable cell type to *bcat-1* disruption. We observed severe degeneration and defasciculation of cholinergic axons/neurites in aged *bcat-1(RNAi)* worms. Consistent with this, several populations of cholinergic cells degenerate in PD, including in the dorsal motor nucleus of the vagus and the nucleus basalis of Meynert (Forno, 1996). In particular, the loss of cholinergic neurons in the pedunculopontine nucleus (PPN) is implicated in gait abnormalities and akinesia in PD (Pahapill and Lozano, 2000), and lesions of the PPN in monkeys produce parkinsonian symptoms (Kojima et al., 1997; Aziz et al., 1998). To our knowledge the present study is the first to link *bcat-1* with cholinergic neurodegeneration. *C. elegans unc-17* and *cha-1* cholinergic synaptic mutants show highly similar motor dysfunction phenotypes to *bcat-1(RNAi)* worms (Sohrabi et al., 2021), further supporting a connection between BCAA metabolism and cholinergic neuron health.

Our results herein suggest that BCAA metabolic defects may contribute to the vulnerability of multiple cell types in PD, providing a potential unifying mechanism of PD neurodegeneration. As demonstrated here, the use of GAIT-GM to uncover previously unknown relationships between genes, metabolites, and functional processes can generate novel testable hypotheses for continued investigation and discovery. Interestingly, we also observed an association between *bcat-1* and reproduction pathways; since *bcat-1* is known to regulate longevity in *C. elegans* (Mansfeld et al., 2015), these findings may suggest another avenue of study that could lend insights into the common trade-off in evolution between reproduction and lifespan. Taken together, our present analyses provide greater understanding of the complex relationship between BCAA metabolism and neuronal health, and our new GAIT-GM tool can be widely applied to biological questions requiring transcriptomic and metabolomic data integration.

## 4 Methods

All method details for the generation of transcriptomics and metabolomics datasets can be found in Mor et al., 2020.

### Transcriptomics

TU3311 uIs60 *(unc-119p::sid-1, unc119p::yfp)* worms were fed either *bcat-1* or control RNAi starting at the L4 stage. On day 5 of adulthood, neurons were isolated and RNA-sequenced (Kaletsky et al., 2016) for five independent collections from each feeding group. Using Galaxy, reads were mapped to the *C. elegans* genome (WS245) using RNA STAR, mapped reads that overlap with gene features were counted using htseq-count (mode = union), and differential gene expression analysis was performed using DESeq2. Raw sequencing reads are available at National Center for Biotechnology Information BioProject PRJNA599166. A total of 3,146 differentially-expressed genes (false discovery rate < 0.05) were selected for the integrated analysis.

### Metabolomics

CF512 *fem-1(hc17); fer-15(b26)* worms were fed either *bcat-1* or control RNAi starting at the L4 stage. On day 5 of adulthood, 6 sets of independent replicates from each feeding group were collected for metabolomics (Mor et al., 2020). Raw data are available at Dryad (https://doi.org/10.5061/dryad.5mkkwh72q). Metabolites were annotated through mummichog (level 3 confidence (Schymanski et al., 2014)), and those that were significantly different (p < 0.05) between the two groups were further examined. If a metabolic feature had multiple putative annotations, the annotation with the least difference between the theoretical and observed mass was retained, and if multiple features were assigned the same putative annotation, the putative annotation with the least difference between the theoretical and observed mass was retained. Adducts and isomers were manually curated for a final list of 110 significantly increased/decreased metabolites selected for the integrated analysis.

#### GAIT-GM tool

GAIT-GM (**G**alaxy **A**nnotation and **I**ntegration **T**ools for **G**enes and **M**etabolites) consists of two major analysis modules (Figure 1, see Supplementary Methods for details) that jointly identify, classify, group and link gene expression and metabolomics data. The *annotation tool* maps gene and metabolites to the KEGG database to enable subsequent pathway-based integration. This novel tool implements a text mining approach to parse metabolite names and match them to KEGG compound IDs. Ambiguous metabolites such as lipids of the same type but different formula’s (i.e. sphingomyelins) are assigned to the same KEGG ID and considered a metabolite class. Gene names are maximally mapped to KEGG through a combination of databases cross-referencing as described (Hernandez-de-Diego, R, 2018). The *integration tool* provides different options for the clustering and summarization of each omic data modality, as well as methods for integrative analysis (see integration workflow examples in Supplementary Figure 3 and Supplementary Figure 4). Metabolites might be grouped by KEGG pathway ID, by metabolite class (i.e. all sphingomyelins), or by abundance patterns (e.g. using the Modulated Modularity Clustering (MMC) tool (Stone and Ayroles, 2009). Similarly, genes can be grouped by pathway and pathway metagenes representing the expression trend in the pathway are computed using an implementation of the PANA algorithm (Ponzoni et al., 2014). Using *correlation tool*, bipartite networks are created to explore the relationship between metabolite and gene expression levels in an unbiased-way and without prior knowledge (Supplementary Figure 4). Alternatively, metabolite-gene associations can be studied by multivariate regression methods (Sparse Partial Least Squares or sPLS; Rohart et al. 2016) where any grouping of explanatory (typically pathway-specific genes or metagenes) and response (i.e. metabolite classes of MMC clusters) variables might be combined. Further information on the GAIT-GM use is provided in the User’s Guide, Tool Input Output, and Methods provided as Supplementary Materials. The tool has been implemented as conda and pypi packages and available at the galaxy toolshed repository (see *Availability of data and materials* section).

#### Integrative analysis of *C. elegans* transcriptomics and metabolomics data

Five biological samples of each experimental condition and omics type were used for integrative analysis. Significant genes and metabolites were uploaded to GAIT-GM and annotated to KEGG pathways using the annotation tool. We used GAIT-GM to analyze the pair-wise correlation between individual metabolites and genes and obtained the bipartite network containing the top 500 metabolite-gene pairs with the largest absolute correlation values. Additionally, we analyzed the relationship between metabolite changes and pathway gene expression changes using the multivariate regression method available in the GAIT-GM. For this, we used KEGG pathway metagenes as explanatory variables and MMC-derived clusters for metabolites as response variables.

##### Gene Ontology

Gene lists were analyzed using gProfiler (Raudvere et al., 2019) with the following settings: *C. elegans* organism, only annotated genes, g:SCS threshold, user threshold 0.05, ENTREZGENE_ACC. REVIGO (Supek et al., 2011) was used to cluster and plot GO terms with a q value < 0.05.

##### Validation Experiments

The following *C. elegans* strain was used for validation experiments: CQ491 vsIs48 [*unc-17p::gfp*]; vIs69 [pCFJ90 *(myo-2p::mCherry + unc-119p::sid-1)*]. This strain was generated by crossing LX929 vsIs48 [*unc-17p::gfp*] with LC108 uIs69 [pCFJ90 *(myo-2p::mCherry + unc-119p::sid-1)*]. Worms were maintained at 20 °C on standard high growth medium (HG) plates seeded with OP50 *E. coli*. Following synchronization by bleaching, worms were transferred at day 1 of adulthood to standard nematode growth medium (NGM) plates supplemented with carbenicillin and isopropyl β-D-1-thiogalactopyranoside (IPTG), seeded with HT115 RNAi *E. coli* (either *bcat-1* or L4440 empty vector control). RNAi plates were pre-induced with 0.1 M IPTG at 1 hour before transfer, and worms were transferred onto fresh RNAi plates every 2-3 days. On day 8, animals were mounted on 2% agarose pads in M9 and sodium azide for cholinergic neuron imaging. Images were captured on a Nikon Eclipse Ti inverted microscope and processed using Nikon NIS elements software. Neurite morphology was scored as normal, moderately degenerated, or severely degenerated at the anterior portion of the ventral and dorsal nerve cords, as well as the neurites projecting from cholinergic neurons in the head ganglia. At least 15 worms were imaged per condition in each replicate.

## Supporting information

Supplementary_User_Guide

Supplementary_Tool_Input_Output

## Declarations

### Ethics approval and consent to participate

Not applicable.

### Consent for publication

Not applicable.

### Availability of data and materials

GAIT-GM scripts, test data, Galaxy xmls are available at https://github.com/secimTools/gait-gm, as a PyPi repository (https://pypi.org/project/gait-gm/), and as a bioconda package (https://anaconda.org/bioconda/gait-gm). A detailed Galaxy User Guide providing step-by-step instructions for running each tool in Galaxy is included as Supplementary Material. All tools are deposited in the Galaxy ToolShed for download and installation (https://toolshed.g2.bx.psu.edu/view/malex/gait_gm/ec9ee8edb84d). Raw RNAseq reads are available at National Center for Biotechnology Information BioProject PRJNA599166 and raw metabolomics data are available at Dryad (https://doi.org/10.5061/dryad.5mkkwh72q).

### Authors’ contributions

DEM, AMM, RK, CTM, VK, GWM, AC, and LMM designed the research; FH coded and implemented the GAIT_GM tool; AMM wrote GAIT-GM documentation and tested the application; OM contributed to GAIT-GM implementation; LMM and AC supervised GAIT-GM designe and implementation. DEM, AMM, RK, VK performed research and analyzed the data; DEM, AC, and LMM wrote the paper.

## Acknowledgements

We acknowledge the support of HiPerGator the University of Florida High performance Computing platform. In particular, the UFL Galaxy instance. We thank the C. elegans Genetics Center for strains (P40 OD010440), and the Confocal Imaging Facility at Princeton University.

