## Supplementary_Figures for "GAIT-GM integrative cross-omics analyses reveal cholinergic defects in a *C. elegans* model of Parkinson’s disease"

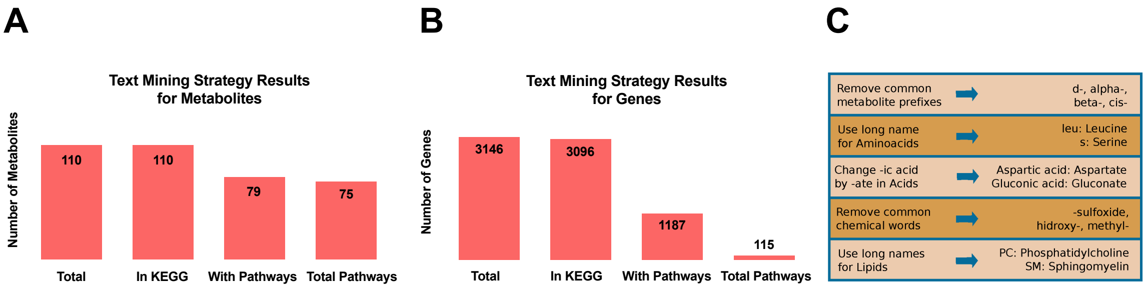


**D**

| Metabolite | Name in KEGG | KEGG ID | Similarity | Tie | Selected | Pathway ID | Pathway Name |
|---|---|---|---|---|---|---|---|
| Ferricyanide | Ferricyanide | cpd:C00324 | 1 | No | Yes | NA | NA |
| Thiamin diphosphate | Thiamin diphosphate | cpd:C00068 | 1 | No | Yes | path:cel00010 | Glycolysis / Gluconeogenesis |
| Thiamin diphosphate | Thiamin diphosphate | cpd:C00068 | 1 | No | Yes | path:cel00020 | Citrate cycle (TCA cycle) |
| Thiamin diphosphate | Thiamin diphosphate | cpd:C00068 | 1 | No | Yes | path:cel00280 | Valine, leucine and isoleucine degradation |
| Thiamin diphosphate | Thiamin diphosphate | cpd:C00068 | 1 | No | Yes | path:cel00620 | Pyruvate metabolism |
| Thiamin diphosphate | Thiamin diphosphate | cpd:C00068 | 1 | No | Yes | path:cel00640 | Propanoate metabolism |
| Thiamin diphosphate | Thiamin diphosphate | cpd:C00068 | 1 | No | Yes | path:cel00730 | Thiamine metabolism |
| Thiamin diphosphate | Thiamin diphosphate | cpd:C00068 | 1 | No | Yes | path:cel01100 | Metabolic pathways |
| Thiamin diphosphate | Thiamin diphosphate | cpd:C00068 | 1 | No | Yes | path:cel01240 | Biosynthesis of cofactors |
| 5-Formiminotetrahydrofolate | 5-Formiminotetrahydrofolate | cpd:C00664 | 1 | No | Yes | path:cel00670 | One carbon pool by folate |
| 5-Formiminotetrahydrofolate | 5-Formiminotetrahydrofolate | cpd:C00664 | 1 | No | Yes | path:cel01100 | Metabolic pathways |
| Tryptophan | L-Tryptophan | cpd:C00078 | 0.9 | No | Yes | path:cel00260 | Glycine, serine and threonine metabolism |
| Tryptophan | L-Tryptophan | cpd:C00078 | 0.9 | No | Yes | path:cel00380 | Tryptophan metabolism |
| Tryptophan | L-Tryptophan | cpd:C00078 | 0.9 | No | Yes | path:cel00400 | Phenylalanine, tyrosine and tryptophan biosynthesis |
| Tryptophan | L-Tryptophan | cpd:C00078 | 0.9 | No | Yes | path:cel00970 | Aminoacyl-tRNA biosynthesis |
| Tryptophan | L-Tryptophan | cpd:C00078 | 0.9 | No | Yes | path:cel01100 | Metabolic pathways |
| Tryptophan | L-Tryptophan | cpd:C00078 | 0.9 | No | Yes | path:cel01210 | 2-Oxocarboxylic acid metabolism |
| Tryptophan | L-Tryptophan | cpd:C00078 | 0.9 | No | Yes | path:cel01230 | Biosynthesis of amino acids |
| Tryptophan | L-Tryptophan | cpd:C00078 | 0.9 | No | Yes | path:cel01240 | Biosynthesis of cofactors |
| Tryptophan | L-Tryptophan | cpd:C00078 | 0.9 | No | Yes | path:cel04361 | Axon regeneration |

**Supplementary Figure 1: Annotation Module applied to the *C. elegans* data.** Metabolite data present a very heterogeneous structure in terms of naming. A set of text mining rules were developed to match the maximum number of terms possible to the unique nomenclature of the KEGG database. Our novel text mining approach maps 100% of metabolites to KEGG nomenclature **(A)** and 98% of genes **(B)**. Note that some of these terms do not have associated KEGG Pathways. **(C)** As abbreviations of terms are common in this kind of data, one solution to match as many terms as possible is to find out the scientific name for each term. Also, removing common chemical words, prefixes and suffixes can improve the percentage of matched terms (details in Supplementary Methods). **(D)** Example of the output of this Annotation Module, where the name in KEGG and the KEGG ID is returned for each input metabolite name, accompanied by the similarity score for matching to KEGG, information about the ties (Tie = Yes indicates cases with the same similarity) and if those metabolites have been selected by the algorithm for downstream integration.

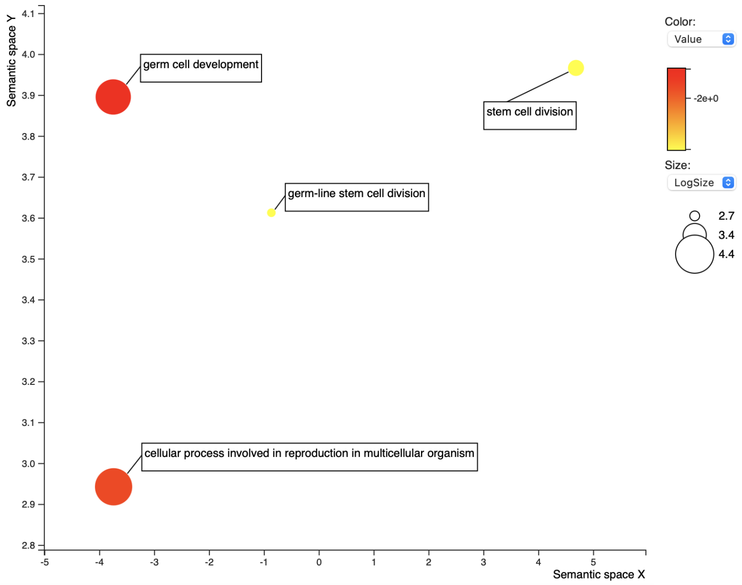


**Supplementary Figure 2:** Gene Ontology analysis visualized by REVIGO for the second largest cluster from **Figure 3A** points to functions related to reproduction.

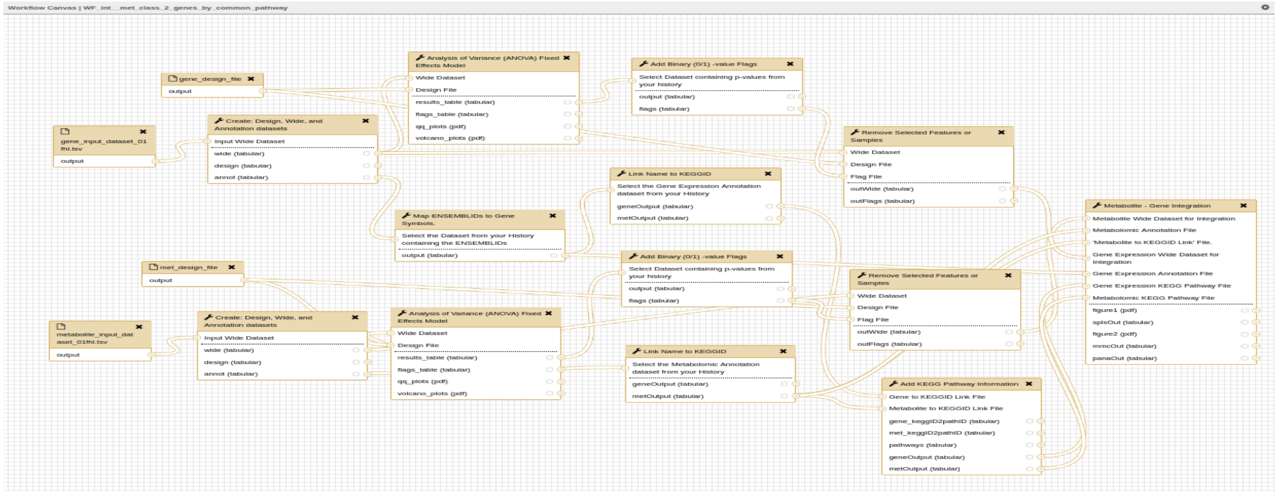


**Supplementary Figure 3:** Example of a Metabolite – Gene Integration Galaxy workflow. Starting with data files and feature identification information (e.g. m/z ratios or retention times), the 'WF_int_met_class_2-genes_by_common_pathway' Galaxy workflow will create wide format datasets, design files, identify the genes and metabolites of interest by ANOVA, annotate the genes and metabolites via KEGG and integrate the gene expression and metabolite data by modeling metabolite classes as a function of the genes in the metabolite pathways. We recommend that both gene expression and metabolite datasets be reduced to reflect a common biological hypothesis before running this tool.

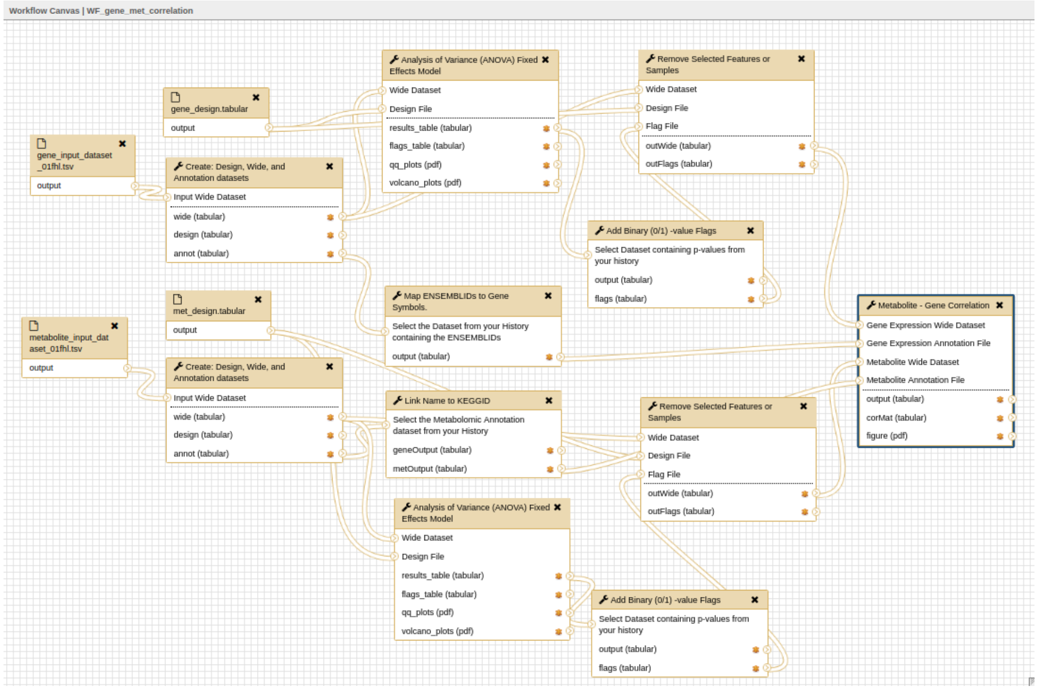


**Supplementary Figure 4:** Metabolite – Gene correlation Galaxy workflow. The workflow starts with the input of gene expression and metabolite abundance data. Metabolite data are subset by class (i.e. using the 'Name_in_KEGG' column generated from the 'Link Name to KEGGID' tool) and only the genes found in the common metabolite pathways are selected for the integration step, here as an unbiased correlation between pairs of genes and metabolites that share a common pathway. The 'WF_int_met_2_metagene.ga' workflow contains the same tools as described above but the options chosen in the 'Metabolite – Gene Integration' tool are different. In this case, the options select model metabolite classes as a function of metagenes where the gene expression data is reduced in scope by implementation of PANA. To include similarly behaving metabolites without regard to identification or type, the 'WF_int_MMC_2_metagene.ga' Galaxy workflow options implement the MMC tool to estimate modules that are modeled as a function of metagenes The 'WF_gene_met_correlation' Galaxy workflow (Supplementary Figure 4) creates wide format datasets, design files, identifies the genes and metabolites of interest by ANOVA, annotates the genes and metabolites via KEGG and performs a correlation analysis between significant genes and metabolites to generate a table of correlation coefficients. P-values for the correlation coefficients are calculated by simulating individual gene and metabolite datasets 1000 times using a normal distribution with means and standard deviations generated from the data. Sample size reflects the input datasets. Correlations are calculated on the simulated data. Correlations must be higher/lower than 95% of the randomly simulated values to be considered significant.
