## Supplementary_User_Guide for "GAIT-GM integrative cross-omics analyses reveal cholinergic defects in a *C. elegans* model of Parkinson’s disease"

### GAIT-GM for Integrative Cross-Omics Analyses
### Galaxy Interface User Guide

GAIT-GM scripts, test data, Galaxy xmls are available at https://github.com/secimTools/gait-gm, as a PyPi repository (https://pypi.org/project/gait-gm/ ), and as a bioconda package (https://anaconda.org/bioconda/gait-gm ). All tools are deposited in the Galaxy ToolShed for download and installation (https://toolshed.g2.bx.psu.edu/view/malex/gait_gm/ec9ee8edb84d ).

The SECIMTools User Guide (Kirpich et al. 2018) describes in detail the structure and format of the input datasets. In brief, wide format data files contain feature measurements for each sample where samples are in columns and features are in rows. These files must contain a column with a unique identifier for each row (feature). Design files are used to relate the samples in the wide format data to sample characteristics (e.g. condition, sex, etc). The sampleID columns in the wide format data files must match the sample names in the 'sampleID' column of the design file. Annotation datasets are wide format data files that relate features to specific feature descriptors (e.g. m/z ratio, compound name, etc.).

**Create: Design, Wide, and Annotation datasets from an Input wide dataset**

This tool takes a single file containing both feature data (e.g. gene or metabolite expression values) and annotation information (e.g. m/z ratio, compound name) and generates the following three files; (1) a wide dataset containing a unique row identifier and the expression values, (2) a wide annotation file with the unique row identifier and any non-data descriptor columns, and (3) a design file with a single column called 'sampleID' with the name of the columns containing the expression data.

If the input dataset does not already contain a column with a unique identifier, the tool will create one. The user can specify a prefix for the unique identifier column (e.g. 'met' for metabolite data). In cases where the input dataset contains a numeric identifier, the tool will append a user-specified prefix or, if no prefix is specified, an underbar. Since the user specifies which columns contain expression values, the resulting wide dataset contains only these data columns and the unique row identifier column. Columns not specified as containing expression values are output into the annotation dataset. The resulting design file template contains a single column called 'sampleID' that contains the names of the user-specified samples in the input data file. The design file can be modified by the user to include additional metadata columns.

Select the **Input Wide Dataset** from the drop-down menu.

Does your Input Wide Dataset have a **unique FeatureID column**?

--If yes, select **Yes.**
--Input the name of the column in your Input Wide Dataset that contains **Unique FeatureIDs**.

Are your unique FeatureIDs ONLY numbers?
--If yes, select **Yes** and
--enter a **prefix to use**, if desired. If no prefix is entered, the tools will prepend an underbar to each numeric FeatureID.
--If no, select **No**.

--If no, select **No**. The tool will generate a unique FeatureID.
--Enter a **prefix to use** when creating the unique identifier. If no prefix is entered, the tool-generated unique FeatureID will be an underbar followed by a number.

Specify which **Sample Columns** contain expression data.
--Enter the numbers (1-based) of the columns in your Wide Dataset that contain expression values. For example, if your expression values are in columns 2-4 then enter: 2,3,4
Columns not specified are treated as annotation descriptor columns. NOTE: annotation descriptor columns are expected to be LEFT of all columns containing expression values.

Click **Execute**.

**Output:**

(1) **Wide TSV dataset**: contains the unique FeatureID column and all user-specified expression data columns.

(2) **Annotation TSV file**: contains the unique Feature ID column and any non-sample descriptor columns.

(3) **Design TSV file**: contains a column called 'sampleID' with the column headers from the input dataset that were selected in step 3. This file can be modified by the user to include additional sample meta-data.

#### Map ENSEMBL IDs to Gene Symbols

This tool takes an annotation data file containing unique FeatureIDs and Ensembl IDs and adds gene symbols. The link from the Ensembl IDs to gene symbols is made using mygene (https://mygene.info/). The tool adds the following columns to the input annotation data file: *GeneSymbol, Score, Selected* and *Tie.*

The *GeneSymbol* column contains the short-form abbreviation for the gene. The *Score* column contains a value generated by mygene indicating how well the Ensembl ID matched the returned gene symbol(s) (Xin, et al. 2016). For cases where an Ensembl ID uniquely matches to a gene symbol, the Selected column = 'Yes". For cases where an Ensembl ID matches to more than one gene symbol, the Selected column = 'Yes' for the gene symbol with the best Score value. If there is a tie, the alphabetically first gene symbol is selected and the Tie column = 'Yes'. We note that FeatureID may not be unique in the resulting output dataset.

1. Select the **species** from which the Ensembl IDs are derived.
   Options include *Homo sapiens, Mus musculus, Rattus norvegicus, Drosophila melanogaster, Arabidopsis thaliana, Sacchoromyces cerevisiae, and Escherichia coli.*

2. Select the **Dataset** containing the Ensembl IDs from your History.

3. Enter the name of the column in your **dataset** that contains the unique **FeatureIDs**.

4. Enter the name of the column in your **dataset** that contains the **Ensembl IDs**.

5. Click **Execute**.

**Output:**

(1) A TSV file containing the data in the input annotation dataset along with the following columns:

**--GeneSymbol:** the Gene Symbol associated with a particular Ensembl ID

**--Score:** a value, generated by mygene, that represents how well the Ensembl ID matched the returned GeneSymbol

**--Selected:** 'Yes' when an Ensembl ID uniquely matches a Gene Symbol or 'Yes' for the best Gene Symbol match when an Ensembl ID matches > 1 gene symbol.

--**Tie:** 'Yes' when an Ensembl ID matches more than one Gene Symbol and all gene symbols have the same Score value.

**<u>Link Name to KEGGID</u>**

This tool takes as input an annotation dataset containing either metabolite compound names or gene symbols and links them to KEGG identifiers (KEGGIDs, KEGG identifiers are unique identifiers for each KEGG object and are also KEGG database entry identifiers) creating either a (a) *Gene to KEGGID Link File* or a (b) *Metabolite to KEGGID Link File*.

For gene symbols, the tool is designed to take the output from the 'Map ENSEMBLIDs to Gene Symbols' tool as input. Therefore, if your gene annotation input dataset contains a column called 'Selected', the tool will link GeneSymbols to KEGGIDs where Selected = 'Yes'. Input files without a column called 'Selected' must have a column of unique FeatureIDs.

For metabolite compounds, the tools links user-specified metabolite names to KEGGIDs by identifying the best match using the following procedure: Common metabolite prefixes are removed (cis-, trans-, d- , l- , (s)-, alpha-, beta-, alpha, beta, alpha-d-, beta-d-, alpha-l-, beta-l-, l-beta-, l-alpha-, d-beta-, d-alpha-). If the metabolite name given is an acid, then the name is modified to the conjugate base by replacing "ic acid", "icacid" or "ic_acid" with "ate". If amino acids are given in 1-letter or 3-letter abbreviations, names are modified to the full amino acid name. The following commonly used lipid abbreviations are modified to reflect the full names (SM = sphingomyelin, lysopc =

lysophosphatidylcholine, PC = phosphatidylcholine, PE = phosphatidylethanolamine and LysoPE = lysophosphatidylethanolamine). Similarly, abbreviations for other commonly assayed metabolites are modified to reflect the full names (cit = citrate, orn = ornithine, thyr = thyroxine and boc = butoxycarbonyl). The code allows the addition of more synonyms. The user-specified metabolite names are retained in the output dataset for comparisons with the KEGG database.
Each parsed metabolite name is compared to metabolite names in KEGG. The best match in KEGG based on similarity score is returned. The similarity score ('Similarity' column) is based on the longest contiguous matching subsequence that does not contain 'junk' elements where 'junk' elements are defined as duplicates making up more than 1% of a sequence with minimum length of 200 (python SequenceMatcher class from difflib)

Selected = 'Yes' for the match with the highest similarity score.

For metabolite names where the best match is tied with at least one other compound in KEGG, all matches are returned. A tie is determined as follows: if the Score is greater than 95% for 2 or more matches in the metabolite name then: 1) the Tie column = 'Yes' and a warning message will appear 2) the Selected column is sorted alphabetically on the Name_in_KEGG column. Note that the user-specified FeatureID and MetaboliteName may not be unique in the resulting output dataset.

1. Select the **species** from the drop-down menu.
Options include *Homo sapiens, Mus musculus, Rattus norvegicus, Drosophila melanogaster, Arabidopsis thaliana, Sacchoromyces cerevisiae, and Escherichia coli.*

2. If you have **both gene and metabolite Annotation Files**, select 'Gene Expression + Metabolomic Annotation Files'.

-- Select your gene expression **Annotation Dataset** from the drop-down menu.

-- Enter the name of the column in your gene expression Annotation File with the **unique FeatureIDs**.

-- Enter the name of the column in your gene expression Annotation Dataset that contains **gene symbols** to link to KEGG identifiers.

-- Select your **metabolite expression Annotation Dataset** from the drop-down menu.

-- Enter the name of the column in your metabolite expression Annotation Dataset containing **unique FeatureIDs**.

-- Enter the name of the column in your metabolite Annotation File that contains the **metabolite names** to link to KEGG identifiers.

3. If you want to use **only a gene expression annotation file**, select 'Gene Expression Annotation File'.

-- Select your **gene expression annotation file** from the drop-down menu.

-- Enter the name of the column in your gene expression annotation file containing **unique FeatureIDs**.

-- Enter the name of the column in your gene expression annotation file that contains the gene symbols to use for linking in KEGG.

4. If you want to use a **metabolite Annotation File only**, select 'Metabolomic Annotation File'.
 -- Select your **metabolite Annotation File** from the drop-down menu.
 -- Enter the name of the column in your metabolite Annotation File containing **unique FeatureIDs**.
 -- Enter the name of the column in your metabolite Annotation File that contains the **compound names to linking to KEGG identifiers**.

5. Click **Execute**.

**Output:**
A TSV file containing the following columns in addition to the columns in the input Annotation file (gene or metabolite):

--**Feature_Type:** whether matching was for metabolites (‘Metabolite’) or genes (‘Gene’).

--**Matched:** whether a match in KEGG was found. ‘Yes’/’No’.

--**Name_in_KEGG**: the KEGG name for the match.

--**KEGG_ID:** column containing the KEGG identifier

--**Similarity:** A score value (range 0 to 1) based on the longest contiguous matching sub-sequence that does not contain 'junk' elements where 'junk' elements are defined as duplicates making up more than 1% of a sequence with minimum length of 200 (python SequenceMatcher class from difflib).

--**Tie:** In cases where multiple matches were found, Tie = ‘Yes’ if the similarity score is greater t han 99%.

--**Selected**: For features with multiple matches and different similarity scores, the 'Selected' column = ‘Yes’ for the match with the highest similarity score. For features with multiple matches and the same similarity score, the 'Selected' column = ‘Yes’ based on alphabetical order.

**<u>Add KEGG Pathway Information using KEGG IDs.</u>**

This tool takes a *Gene to KEGG Link File* or a *Metabolomic to KEGG Link File* and adds KEGG pathway names using the KEGG identifiers. The tool was designed to take the output from the previous ‘Link Name to KEGG ID' tool as input (for example, the 'Gene to KEGG ID link File’) but other datasets containing KEGG IDs and can be used as well.

If a '*Gene to KEGG ID link File*’ is given, the tool outputs the following three files;

1) a '*Gene KEGG Pathway File*' containing the unique FeatureID, Feature_Name, Feature_Type and KEGG_ID columns from the input file plus Pathway_ID and Pathway_Name columns containing KEGG pathway identifiers and KEGG pathway names, respectively. If no pathways are linked to the KEGG IDs then these columns return 'NA'.

2) a '*GeneKeggID2PathwayID*' file containing the KEGG gene identifiers and their associated KEGG pathway identifiers.

3) a '*PathwayID2PathwayNames*' file containing all of the KEGG pathway identifiers and their associated KEGG pathway names.

If a '*Metabolite to KEGG ID link File*' is given, the tool outputs the following three files;

1) a ' *Metabolite KEGG Pathway File*' containing the unique FeatureID, Feature_Name, Feature_Type and KEGG_ID columns from the input file plus Pathway_ID and Pathway_Name columns containing KEGG pathway identifiers and KEGG pathway names, respectively. If no pathways are linked to the KEGG IDs then these columns return 'NA'.

2) a '*MetaboliteKeggID2PathwayID*' file containing the KEGG metabolite identifiers and their associated KEGG pathway identifiers.

3) a '*PathwayID2PathwayNames*' file containing all of the KEGG pathway identifiers and their associated KEGG pathway names.

Note: FeatureIDs and KEGGIDs may NOT by unique in the output files.

1. Select your **species** from the drop-down menu.
   Options include *Homo sapiens, Mus musculus, Rattus norvegicus, Drosophila melanogaster, Arabidopsis thaliana, Sacchoromyces cerevisiae, and Escherichia coli.*

2. If you have **both gene and metabolite Annotation Files with KEGGIDs**, select 'Gene Expression + Metabolomic Files with KEGGIDs'.
   -- Select your '***Gene to KEGG ID Link File***' from the drop-down menu.
   -- Enter the name of the column in your '*Gene to KEGG ID Link File*' with the **unique gene FeatureIDs**.
   -- Enter the name of the column in your '*Gene to KEGG ID Link File*' that contains **gene symbols**.
   -- Enter the name of the column in your '*Gene to KEGG ID Link File*' that contains **KEGG gene identifiers**.
   -- Select your '***Metabolite to KEGG ID Link File***' from the drop-down menu.
   -- Enter the name of the column in your '*Metabolite to KEGG ID Link File*' with the **unique FeatureIDs**.
   -- Enter the name of the column in your '*Metabolite to KEGG ID Link File*' that contains **metabolite names**.
   -- Enter the name of the column in your '*Metabolite to KEGG ID Link File*' that contains the **KEGG metabolite identifiers**.

3. If you have **a Gene Annotation File with KEGG IDs**, select 'Gene Expression File with KEGGIDs'.

-- Select your '***Gene to KEGG ID Link File***' from the drop-down menu.
-- Enter the name of the column in your '*Gene to KEGG ID Link File*' with the **unique gene FeatureIDs**.
-- Enter the name of the column in your '*Gene to KEGG ID Link File*' that contains **gene symbols**.
-- Enter the name of the column in your '*Gene to KEGG ID Link File*' that contains **KEGG gene identifiers**.

4. If you have **a Metabolite Annotation File with KEGG IDs**, select 'Metabolomic File with KEGGIDs'.
-- Select your '***Metabolite to KEGG ID Link File***' from the drop-down menu.
-- Enter the name of the column in your ' *Metabolite to KEGG ID Link File*' with the **unique metabolite FeatureIDs**.
-- Enter the name of the column in your ' *Metabolite to KEGG ID Link File*' that contains **metabolite names**.
-- Enter the name of the column in your ' *Metabolite to KEGG ID Link File*' that contains **KEGG metabolite identifiers**.

5. Click **Execute**.

**Output:**

For either metabolite or gene annotation input files, the following 3 output files are generated:

The following files are generated if the '*Gene/Metabolite to KEGG ID Link File*' is selected:

1. **Gene/Metabolite KEGG Pathway File.** TSV file containing following columns:
   UniqueID: unique identifier, numeric
   Feature_Name: gene symbol or metabolite name
   Feature_Type: 'Gene' or 'Metabolite'
   KEGG_ID: KEGG identifier
   Pathway_ID: KEGG pathway identifier, if found
   Pathway_Name: pathway name in KEGG

2. **Gene/MetaboliteKeggID2PathwayID.** Downloaded information from KEGG that contains:
   1. column of KEGG pathway identifiers
   2. column of KEGG identifiers

3. **PathwayID2pathwayNames.** Downloaded information from KEGG that contains:
   1. KEGG pathway identifier
   2. KEGG pathway name

**<u>Add Binary (0/1) P-value Flags</u>**

This tool generates 0/1 indicator variables to identify ('flag') p-values less than a user-specified threshold or less than the default values of 0.1, 0.05 and 0.01. The *flag_threshold* variable is equal to 1 if the p-value is less than or equal to the threshold and equal to 0 if greater than the threshold. You can flag nominal p-values or p-values after correction for multiple testing.

1. Select the **dataset** containing a column of **p-values** you wish to flag from your History.

2. Enter the name of the column in your dataset that contains the **unique FeatureIDs**.

3. Enter the name of the column in your dataset that contains the **p-values**.

4. Enter the **p-value threshold(s)** for flagging. P-values less than the given threshold(s) will be flagged with a 1. If you enter more than one threshold value, separate the values with a comma (no spaces). Default values are 0.1, 0.05, and 0.01.

**Output:**

Two output files are generated by the tool:
**1. Flags File.** A TSV file containing the unique FeatureID columns plus the additional binary indicator columns for whether a p-value was less than the threshold:

--uniqueID
--flag_05: binary indicator flag where a 1 indicates the p-value was less than 0.05
--flag_01: binary indicator flag where a 1 indicates the p-value was less than 0.01
--flag_1: binary indicator flag where a 1 indicates the p-value was less than 0.1

**2. Output File.** A TSV file containing the same columns as the input dataset plus the binary indicator flags.

#### Metabolite – Gene Correlation

The tool performs a correlation analysis between genes (*Gene Expression Wide Dataset*) and metabolites (*Metabolite Wide Dataset*) to generate a table of correlation coefficients. P-values for the correlation coefficients are calculated by simulating gene and metabolite datasets 1000 times using the mean and standard deviation of both datasets.

The tool outputs 2 TSV files and a PDF figure. The 'correlation file' contains gene-metabolite correlation coefficients with p-values less than the user-specified threshold. The tool also outputs the results in matrix format, the ‘correlation matrix file’. The ‘correlation figure’ is a network representation of the top 500 gene-metabolite correlations based on the absolute value of the correlation coefficients.

1. Select the **Gene Expression Wide Dataset** from your History.

2. Enter the name of the column in your Gene Expression Wide Dataset with **unique FeatureIDs**.

3. Select whether you want to use a **Gene Expression Annotation File**. Selecting this option allows the user to chose a column in the annotation file for labeling output files (e.g. gene names).
--if you select **'Yes'**
--Select the **Gene Expression Annotation File** from your History.
--Enter the **name of the column** in your Gene Expression Annotation File to use for labeling.

4. Select the **Metabolite Wide Dataset** from your History.

5. Enter the name of the column in your Metabolite Wide Dataset that contains the **unique FeatureIDs**.

6. Select whether you want to use a **Metabolite Annotation File**. Selecting this option allows the user to chose a column in the annotation file for labeling output files (e.g. compound names).
  --if you select **'Yes'**
    --Select the **Metabolite Annotation File** from your History.
    --Enter the **name of the column** in your Metabolite Annotation File to use for labeling.

7. Select a **correlation method**. Options include Pearson (Pearson's standard correlation coefficient, default value), Spearman (Spearman's rank correlation) or Kendall (Kendall's Tau correlation coefficient).

8. Enter a **p-value threshold**. This value is used to filter the results in the '*Correlation File*' to only those correlations whose p-value is less than this threshold. Default 0.05.

8. Click **Execute**.

**Output:**

Two data files and a PDF are generated by the tool:

**1. Correlation File.** TSV file, sorted by the absolute value of the correlation coefficient, that contains gene-metabolite correlation coefficients with p-values less than the specified threshold.
  -- Gene: gene identifier column
  --Metabolite: metabolite identifier column
  --correlation: correlation coefficients
  --p-value: p-value generated from the mean and standard deviation of the input gene and metabolite wide datasets

**2. Correlation Matrix.** TSV file of the correlation coefficients in matrix format

**3. Correlation Figure.** PDF figure containing a network representation of the top 500 gene-metabolite correlations based on the absolute number of the correlation coefficient. Max number of correlation coefficients is 500.

**Metabolite – Gene integration**

Note: The parameters you select are data dependent.

This tool carries out an integrated analysis of metabolite and gene expression data. Metabolite data are the dependent variable and genes are the explanatory variable. The tools allows for several combinations of metabolite and gene models. A note of caution: an integrated analysis of metabolite and gene expression dataset with no filtering will be challenging to interpret using this tool. We recommend that both datasets be reduced to reflect a common biological hypothesis before running the tool.

Options for subsetting metabolite expression data include:
(1) by metabolite class (uses the 'Name_in_KEGG' column in the output generated from the 'Link Name to KEGG ID' tool),
(2) by MMC (Modulated Modularity Clustering) pattern (Stone and Auroles, 2009, Kirpich et al. ) to estimate similar underlying modules) and
(3) by both metabolite class and MMC.

Options for subsetting of gene expression data include:
(1) no reduction,
(2) a custom TSV file containing specific genes of interest,
(3) genes linked to each metabolite class through common KEGG pathways (uses the 'Gene and Metabolite KEGG Pathway files' generated from the 'Add KEGG Pathway Information' tool), and
(4) metagenes from PANA (PAthway Network Analysis from gene expression data, Ponzoni et al. 2014).

1) **Classes of metabolites can be modeled as a function of metagenes.**
2) **Classes of metabolites can be modeled as a function of a set of individual genes.**
3) **Unbiased clusters of metabolites can be modeled as a function of metagenes**
4) **Unbiased clusters of metabolites can be modeled as a set of individual genes**

The tool executes a partial least squares regression with variable selection (sparse PLS, sPLS) as implemented in the 'mixOmics' package ( Rohart et al. 2017). The sPLS function is run in 'classic mode' (http://mixomics.org/methods/spls/) with the number of components included in the model set to 2. In addition, the user selects the number of variables (genes) for each component to use in model construction.

1. Select a **Metabolite Wide Dataset** for Integration from your History.

2. Enter the name of the column in your Metabolite Wide Dataset that contains the **unique FeatureIDs**.

3. Optional: Select a **Metabolite Annotation File**. Selecting this option allows the user to chose a column in the Annotation File for labeling output files.

– if you select **'Yes'**
  – Select the **Metabolite Annotation File** from your History.
  – Enter the **name of the column** in your Metabolite Annotation File to use for labeling.

4. Select an option for modeling/subsetting the Metabolite Data.

--(1) **'By metabolite class'** To use a predefined grouping of metabolites.

--all of the metabolites in each class (1,..j) will be treated as individual dependent variables in a multivariate regression ($Y_1$,..$Y_{n\_j}$) where j is the number of metabolites in class j.

--Select the '***Metabolite to KEGG ID Link***' File from your history. This file MUST contain a column called 'Name_in_KEGG' that is used to define the metabolite groups.

--(2) '**By MMC pattern**' Run the SECIMTools MMC tool. Each resulting MMC module will be analyzed as a set.

--Select the '**Metabolite KEGG ID Link**' File from your history

--Select the **Design File** to use with your Metabolite KEGG ID Link File. This file can be generated using the 'Create: Design, Wide, and Annotation datasets' tool. Note: at minimum you need a column called 'sampleID' that contains the names of your samples.

--The following SECIMTools MMC options are described fulling in Aroyles citation.
--Lower sigma bound
--Upper sigma bound
--Number of sigma values
--Correlation method

--(3) '**By both metabolite class AND MMC pattern**' Options as described above.

5. Select a **Gene Wide Dataset** for Integration from your History.

6. Enter the name of the column in your Gene Wide Dataset that contains the **unique FeatureIDs**.

7. Optional: Select a **Gene Annotation File**. Selecting this option allows the user to chose a column in the Annotation File for labeling output files.

– if you select **'Yes'**
– Select the **Gene Annotation File** from your History.
– Enter the **name of the column** in your Gene Annotation File to use for labeling (e.g gene names).

8. Select an option for modeling/subsetting the Gene Data.

--(1) **'Include all genes'** No subsetting.
Note: run times may be so long that the process fails. Not recommended.

--(2) '**Upload a custom list containing specific genes of interest**' This list must be a single column containing Gene Symbols for the genes of interest

--Select the **Custom Gene List** from your History

--(3) **Use Metagenes** (PANA)

--Select the '**Gene to KEGGID Link File**' from your History

--Enter the name of of the column in the 'Gene to KEGGID Link File' that contains **gene symbols.**

--Select the '**GeneKeggID2PathwayID**' File from your History. This file contains ALL gene KEGG IDs to Pathway IDs and can be generated from the 'Add KEGG Pathway

Information' tool.

--Choose the **criterion to select components**. Options are:
--single%, percent of variability for a given principle component
--%accum, percent of accumulated variability
--abs.val, absolute value of the variability for a given principle component
--rel.abs, fold variability of total variability divided by rank(X).

--Enter a **variability cut-off value**. Default of 0.23

--Select whether to include **Pathway names** in the output files and figures
--if you select **'Yes'**
--Select the '**GeneKeggID2PathwaNames**' File from your History. This file can be generated from the 'Add KEGG Pathway Information' tool.

9. Enter the **number of genes to keep** for each component in the sPLS analysis.

10. Enter the **correlation threshold**. Correlations less than this value will not be included in output files. Used to make visualization of results manageable. Default value of 0.8.

11. Click **execute**.

**Output:**

**For metabolite reduction by metabolite class and all genes:**
(1) PDF sPLS figure for each metabolite class
(2) sPLS correlation TSV file containing the correlations for each gene-metabolite pair and what metabolite class (subset) the pair locate to.

**For metabolite reduction by MMC:**
The following will be output in addition to files (1) and (2) above:

(3) MMC PDF figure containing unsorted, sorted and sorted-smoothed heatmaps of the variance-covariance matrixes

(4) MMC TSV file containing algorithm summaries in the following columns:
--unique metabolite FeatureID
--Module: module number for each feature calculated by MMC tool
--Entry Index: original order of row names for the input Metabolite Wide Dataset
--Degree: average of the absolute values of the correlations for the given element in a block to other elements within that block
--Average Degree: average values of the degrees computed above across all elements with the given block

**For subsetting genes by generating megagenes using PANA:**
The following additional files will be output:

(5) PANA TSV file containing the associations between gene symbols and KEGG pathways
--Metabolite: metabolite name

--Gene: gene symbol
--Correlation: correlation coefficient
--Subset: subset ( ametabolite class, MMC module, etc.)
