## Supplementary_Methods for "GAIT-GM integrative cross-omics analyses reveal cholinergic defects in a *C. elegans* model of Parkinson’s disease"

### Overview and availability

There are three basic steps for an integrated expression and metabolite analysis 1) Feature annotation 2) Feature selection 3) Integrated analysis. We have developed Galaxy tools that cover the first and third steps of this process. GAIT-GM **G**alaxy **A**nnotation and **I**ntegration **T**ools for **G**enes and **M**etabolites. Here we describe the methods for the tools that we developed to facilitate this process. These tools are deployed in Galaxy with existing tools leveraged in workflows. In Supplementary Material: Results and Workflows we provide examples of integrated omics analysis. All of the tools are wrapped for Galaxy and deposited in PyPi (https://pypi.org/project/gait-gm/) with a corresponding Conda recipe so sophisticated users have access on the command line. Details are provided in the Supplementary Material: User Guide. All code, including the needed Galaxy wrappers is also available on our github (https://github.com/secimTools/gait-gm).

### Annotation

*Linking metabolites and genes to KEGG identifiers – 'Link Name to KEGGID'*

In order to integrate transcriptomics and metabolomics data using common KEGG pathways, each omics data needs to be converted into the identifiers used by the KEGG database. Genes in KEGG are not directly attached to common identifiers such as RefSeq/PubChem identifiers, rather, KEGG uses an independent identifier (a KEGGID). Similarly, metabolite name conventions are diverse and KEGG uses their own compound IDs. We have developed tools to map input gene/metabolite names to the KEGG database. The KEGG identifiers can then be retrieved and used to link to KEGG pathways. This process identifies which genes and metabolites are in shared pathways.

*Resolving gene names*

Gene ID conversion has been a long and general problem of genomics databases. Tools such as DAVID (Dennis et al. 2003) and BridgeDB(van Iersel et al. 2010) address this problem, although limitations exist, such as the number of species covered or the number of items that can be processed at a time. The KEGG Mapper is inconsistent in its naming conventions for different species. To our knowledge there is no general tool that links KEGGIDs for all species. We have adopted the PaintOmics3 procedure for gene mapping (Hernandez-de-Diego et al. 2018). Basically, PaintOmics3 fetches the ID translation information from public databases such as Ensembl, PDB, NCBI Refseq and KEGG, and generates the translation tables and stores them in a MongoDB collections. For example, given a featureID (gene, protein or transcript) for database A, to translate to a valid gene name for database B, first the system retrieves the list of transcripts associated with the feature (if any). Then, for each transcript ID in database A, the equivalent transcript identifier at database B. Finally, to translate back to genes, the system finds the gene name associated to each identified transcript. Although this method has some limitations, mainly due to the fact that intersections between databases are not complete (i.e. some biological entities in database A may not exist in database B), in general terms the percentage of translated features has shown to be high and sufficient enough for pathway analysis purposes (Hernandez-de-Diego et al. 2018).

*Resolving metabolite names*

There are several different IDs that exist for metabolites such as InChIKeynd PubChemID(Kim et al. 2016), If available, an existing tool (Wohlgemuth et al. 2010) can be used to batch convert these ID's to the KEGGID to another. However, despite the availability of IDs, and the knowledge that metabolite names are suboptimal (Kind, Scholz and Fiehn 2009) natural language names are often given to the scientist by the core facility. While we recognize that giving compound names rather than IDs is not a best practice, it is simply the current reality for many using metabolomics technology. An additional complication is that using names for metabolites introduces variability due to different technology providers using different conventions. For example, using *lactate* versus *lactic acid (*the conjugate base of an acid versus the acid) or using *α-ketoglutarate* versus 2-*oxoglutarate*. Moreover, metabolomics platforms usually do not resolve isomeric variants of sugars such as L-glucose and D-glucose, for which a different notation exists in KEGG. At present, common natural language names are not standardized in the chemical community. We map names onto KEGGIDs using a set of rules for the processing of natural language. We took the initial set of rules in PaintOmics3 (Hernandez-de-Diego et al. 2018) and have expanded upon that process to assign metabolite names to the most likely compound in the KEGG database. For each input metabolite, a list of potentially related metabolites based on the similarities in their names is generated as follows: 1) Metabolite names are parsed according to the rules described below. 2) Names are matched to KEGG 3) a similarity score is calculated using the python internal SequenceMatcher class from difflib (https://docs.python.org/2/_sources/library/difflib.rst.txt, module and section author Tim Peters) that returns a measure of the similarity between two strings. Identical names receive a score of 1. 4) The highest similarity score is selected 5) When the best match is tied with at least one other compound in KEGG, all matches are returned. A tie is determined as follows if the similarity score (greater than 95% for 2 or more matches in the metabolite name). A default selection is made, but other possibilities are visible to the scientist and the selection can be easily modified before the next step.

*Text mining rules*

Assigning a KEGG ID, KEGG Name and KEGG Pathway: Metabolites / small molecules are referred to by a large number of names and abbreviations, some of these depend on the database used. In (many) cases an experiment cannot unambiguously identify and name a metabolite, instead only determine a metabolite "class". To assign KEGGIDs to metabolites / metabolite classes identified in an experiment, a set of text mining rules was developed to match experiment names to unique compound names in the KEGG database. This approach maximizes the number of terms in common. After an initial attempt at a full name match, experiment names are simplified using the following procedure. Common metabolite prefixes are removed (cis-, trans-, d- , l- , (s)-, alpha-, beta-, alpha, beta, alpha-d-, beta-d-, alpha-l-, beta-l-, l-beta-, l-alpha-, d-beta-, d-alpha-). If the metabolite name given is an acid, then the name is modified to the conjugate base by replacing "ic acid", "icacid" or "ic_acid" with "ate". If amino acids are given in 1-letter or 3-letter abbreviations, names are modified to the full amino acid name. The following commonly used lipid abbreviations are modified to reflect the full names (SM = sphingomyelin, lysopc = lysophosphatidylcholine, PC = phosphatidylcholine, PE = phosphatidylethanolamine and LysoPE = lysophosphatidylethanolamine). Similarly, abbreviations for other commonly assayed metabolites are modified to reflect the full names (cit = citrate, orn = ornithine, thyr = thyroxine and boc = butoxycarbonyl). Each parsed metabolite name is compared to metabolite names in KEGG. The best match in KEGG is identified based on the longest contiguous matching subsequence that does not contain 'junk' elements where 'junk' elements are defined as duplicates making up more than 1% of a sequence with minimum length of 200 (python SequenceMatcher class from difflib). Note that the experimental metabolite names and information about any ties are retained in the output dataset for manual curation. This procedure increases the number of matches between experimental metabolites and metabolites in KEGG.

*Features to Pathways – 'Add KEGGID to Pathway Information'*

Linking annotations form the data to a common identifier such as a KEGG identifier is the first step in an integrated the analysis. The next step is to link the KEGG identifiers to Pathways, which is a straight-forward parsing KEGG pathway files. Here it is important to keep in mind that a single feature (metabolite or gene) may be in multiple pathways.

***Selection and filtering***

We recommend performing a selection of metabolites/genes prior to the integration task. For example, differential expression can be used to filter metabolites/genes changing between treatments (Supplementary Materials: Results and Workflows). Feature selection can be performed with any Galaxy tool that implements differential expression analysis. For example, for gene expression, the Galaxy implementations of edgeR or DEseq2 can be used. For metabolomics, we recommend using SECIMTools (Kirpich 2018). These tools return lists of features (genes or metabolites) with associated p or q values that can be used for threshold-filtering.

### Integration

We have designed a framework for the integration of metabolomics with gene expression data that builds on four biologically inspired notions and the principle of analysis flexibility (Figure). The first notion is that gene expression regulates metabolite levels and this regulatory relationship can be discovered. The second notion is that genes/metabolites belonging to the same pathway are more likely to be engaged in regulatory relationships. The third notion is that metabolites/genes that belong to the same "class" (i.e. all sphingomyelins; have co-expression patterns) have a shared latent structure. Fourth, profiles of co-expressed genes can be considered as a proxy for an underlying regulatory element that may contributes to metabolic regulation of a class of metabolites. The principle of flexibility implies that multiple combinations or subsets of genes and metabolites can be integrated to answer different research questions. GAIT-GM consists, therefore, in a number of functions that implement these concepts.

*Defining metabolite subsets*

Metabolites can be analyzed as a function of gene expression as a whole data matrix, or as subsets of metabolites. Subsets can be created either by their annotation label or by their measured levels across samples. Annotation labels refer to the metabolite ID in the KEGG database, which can be assigned to multiple compounds of the input dataset. For example, when lipidomics data are processed, multiple compounds will be mapped to the KEGG ID sphingomyelin or ceramide, and hence these compounds are considered to be part of the same metabolite class. Metabolite groups can also be created based on their measurements across samples using a clustering technique. Here we use the SECIMTools (Kirpich et al. 2018) implementation of MMC (Stone and Ayroles 2009) for unbiased clustering or metabolites. Note that subsets can be created by combining the annotation class with the MMC cluster or by a user defined knowledge base.

*From genes in pathways to metagenes.*

Gene expression can be used a whole data matrix or genes can be selected that belong to specific pathways. An additional possibility is to concentrate pathway gene expression information in one or few metagenes that capture the variability pattern or "activity" of the pathway across samples. We have implemented the pathway metagene computation method described in Ponzoni et al 2014. This typically reduces the gene expression dataset to as many metagenes as annotated pathways.

*Metabolite-Expression integration statistics.*

*Gene-metabolite correlation*

A simple **correlation measure** is available to calculate correlations between metabolite abundance and gene expression for all possible gene-metabolite pairs used as input (Figure 1). The tool then selects the top 500 correlation pairs to display data as an interaction network, that [illegible]

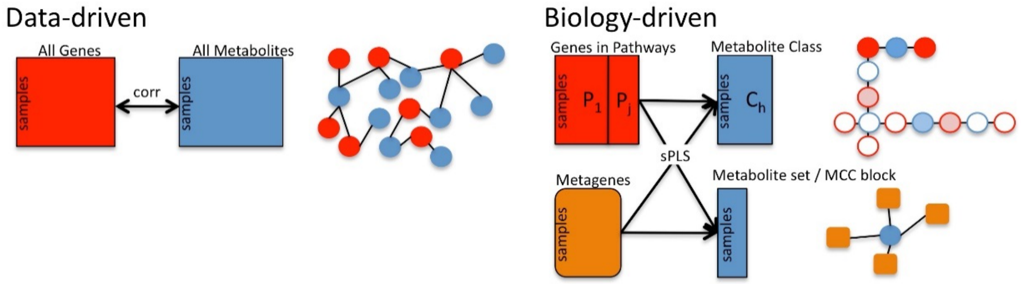


*Figure 1 The gene-metabolite correlation tool allows an unbiased comparison of metabolite abundance to gene expression. The gene-metabolite integration tools uses structured subsets to facilitate the modeling of metabolites as a function of gene expression*

[illegible] at [illegible] o-variation relationships between genes and metabolites.

Given the high number of correlations calculated here, it is important to estimate the potential for spurious association, GAIT does a simulation test, where the mean and variance of each compound/gene is assumed to be normal and this used to generate observations at random for all features used. The simulated data are then processed identically to the observed data and the distribution of the correlation coefficients are calculated. The simulation is performed 1000 times, and for each possible gene/metabolite pair, the frequency of random correlations above the correlation obtained with the original data is taken as p.value (Figure 2). Multiple testing may be adjusted for by several different already existing tools in Galaxy such as the 'Multiple Testing Adjustment (MTA)' tool implemented in SECIMTools (Kirpich et al. 2018).

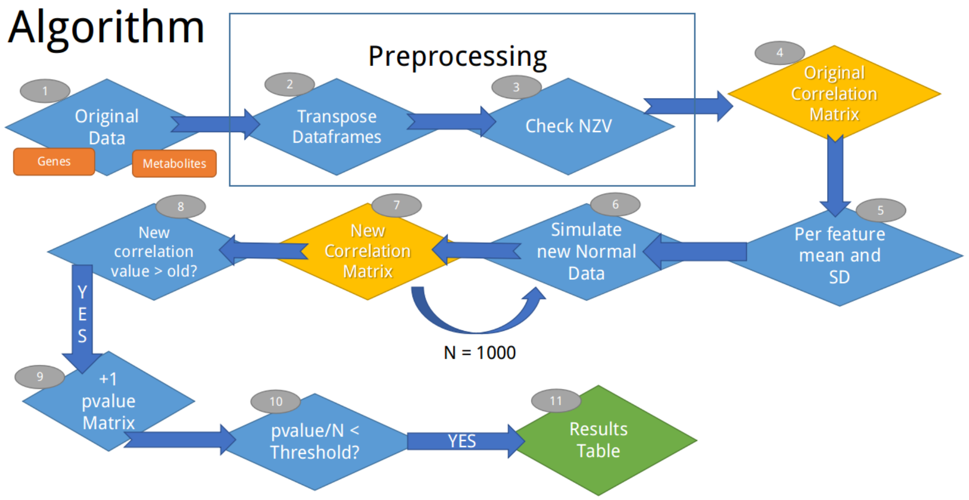


*Figure 2. For each gene-metabolite pair a correlation is calculated. For each gene and each metabolite, the mean and std are estimated from the data. Using those numbers, random values are drawn from a normal distribution and correlations are calculated using those random values. If the simulated value is more extreme than the value from the data, a counter is incremented. A p-value is estimated as the number more exptreme/ the total number of simulations. If this is smaller than the user provided threshold the results are considered potentially significant and retained.*

*Integration through multivariate statistics.*

We implement **Sparse Partial Least Squares** (sPLS) from the mixomics package (Rohart et. al. 2016) as a method to explain metabolite level changes as a function of gene expression. In this approach, Gene Expression is the explanatory variable (X) and Metabolite levels is the response variable (Y). This statistical method can be applied to multiple combinations of Gene Expression and Metabolite matrices to create a highly flexible analysis framework where several regulation hypotheses can be tested. We next explain some of these combinations:

a) <u>Metabolite class vs. genes in associated pathways</u>. A group of metabolites in the input data that map to the same compound ID in the KEGG database are considered a class and represent a single feature that might be present in one or several KEGG pathways. A pertinent question in this case is if all the metabolites in the same class are regulated in the same way or subsets of metabolites associate to different sets of genes. The sPLS model in this case permits to answer this question. By using as explanatory variables only genes in the pathway where the metabolite is annotated, we increase the chance of identifying meaningful associations.

b) <u>Metabolite class vs. pathway metagenes.</u> Annotation of metabolites to metabolic and signaling pathways is far from complete and we might want of explore regulatory relationships beyond the pathways where metabolites are currently annotated. Using in the case the whole gene expression space might introduce too much noise. As sensible approach is to summary pathway activity as the combination of the expression profiles of genes in the pathway and use these pathway metagenes as explanatory variables to construct the sPLS models. When the response variable is a metabolite class this approach addresses the question of how the pathway activity network contributes to the regulation of the metabolites in the same class.
